# Tree cavities as attractors: an experimental test of the effects of woodpecker legacy on local bird activity and species richness

**DOI:** 10.64898/2026.11.05.764588

**Authors:** Alicja Wolska, Marcin Trybała, Michał Ciach

## Abstract

Tree cavities, whether excavated by birds or formed through wood decay, are key structural features of forest ecosystems. They are critical resources for secondary cavity users and influence breeding bird assemblages by affecting the occurrence and abundance of certain species. However, the effect of an individual cavity on fine-scale bird activity, independently of the surrounding habitat context, remains poorly understood. Here, we experimentally tested the effect of cavity presence on bird activity in a coniferous forest in southern Poland. We installed pairs of standardized, debarked snags, each consisting of one snag with an artificial entrance resembling a woodpecker-excavated cavity and one control snag without a cavity entrance. Bird presence and activity were monitored using camera traps and vegetation structure was quantified around each snag. The presence of a cavity entrance had a positive effect on visitation rates and species richness. Snags with cavity entrances received 114% more bird visits than control snags (4.5 [95% CI: 3.1–6.5] vs 2.1 [1.4–3.2]) and were visited by 48% more species (2.0 [1.5–2.6] vs 1.3 [0.9–1.9]). In addition, greater canopy closure around snags was associated with higher visitation rates and species richness. Our study demonstrates that a single cavity can act as a fine-scale attractor for forest birds. These findings highlight the role of cavities and cavity-bearing trees in shaping spatially heterogeneous patterns of fine-scale habitat use by birds.

**Highlights:**

- Cavity presence acts as a fine-scale attractor for forest birds
- Artificial cavities increase visit frequency and bird species richness
- Cavity-associated species dominate among birds attracted to cavities
- Cavity presence influences the behaviour of birds visiting snags

## 1. Introduction

The concept of keystone species posits that certain organisms exert effects on ecosystem structure and functioning that are disproportionate to what would be expected from their abundance or biomass (Paine, 1966). This well-established ecological concept has important implications for ecology and conservation, as the presence or loss of keystone species can influence community composition, species interactions and ecosystem processes. Consequently, identifying and protecting keystone species may be important for maintaining ecosystem functioning and biodiversity (Mills et al., 1993). In forest ecosystems, primary cavity nesters, particularly woodpeckers (family Picidae), are often considered keystone species (Virkkala, 2006; Remm & Lõhmus, 2011), as the cavities they excavate provide essential resources for a wide range of taxa (Cockle et al., 2019). Woodpecker-excavated cavities are commonly used by secondary cavity users, particularly birds (Martin & Eadie, 1999; Wesołowski & Martin, 2018). Tree cavities provide suitable nesting conditions, including a relatively stable microclimate that buffers occupants from ambient environmental conditions (Wiebe, 2001; Mersten-Katz et al., 2012; Maziarz et al., 2017). By providing nesting and roosting sites, cavities influence species composition and abundance, thereby shaping bird assemblages (Drever et al., 2008; Cockle et al., 2011; Van Der Hoek et al., 2020). The ecological relationships among cavity excavators, cavities and secondary cavity users form a network known as a nest web (Martin & Eadie, 1999; Martin et al., 2004).

Forest management reduces the availability of cavity-bearing trees, potentially affecting the occurrence and abundance of some bird species and, consequently, ecosystem functioning (Remm & Lõhmus, 2011; Edworthy & Martin, 2014; Walankiewicz et al., 2014; Andersson et al., 2018). To compensate for this loss of nesting opportunities, nest boxes are widely used worldwide to support cavity-dependent species (Dulisz et al., 2022; Thompson et al., 2023). In addition, several methods have been developed to create cavities or accelerate their formation in trees (Carey & Sanderson, 1981; Copeyon et al., 1991; Griffiths et al., 2018; Ellis et al., 2022). These conservation approaches primarily treat cavities as resources that provide potential nesting or roosting sites. However, the ecological influence of a cavity may extend beyond its direct use as a nesting or roosting site, as birds may inspect and repeatedly visit cavities and their immediate surroundings for various purposes (Forsman & Thomson, 2008; Tolvanen et al., 2018). Given that cavities can be abundant under natural conditions (Przepióra & Ciach, 2023), such visits may also have implications for ecological processes within forest ecosystems.

Although the ecological importance of cavities is well established, most studies have examined their role in structuring bird communities at the stand or landscape scale (Martin & Eadie, 1999; Remm & Lõhmus, 2011; Trzcinski et al., 2022). In contrast, it remains unclear whether the mere presence of an individual cavity can alter bird activity and species composition at a local scale. Assessing cavity attractiveness under natural conditions is challenging because natural cavities vary in their quality, tree species, stage of decay, height, dimensions and orientation (Martin et al., 2004; Remm et al., 2006). Moreover, cavities that are currently or have previously been used by birds or other vertebrates may provide secondary cavity users with additional sources of social information, such as faeces, nest remains and other organic material (Mazgajski, 2007; Podofillini et al., 2018), potentially influencing their behaviour. Consequently, the independent effect of cavity presence on bird behaviour is difficult to disentangle from the characteristics of cavity-bearing trees and the surrounding habitat context.

In this study, we assessed the effects of a single cavity on local bird activity, species richness and behaviour. To isolate the effect of cavity presence as a structural feature while minimising the influence of environmental heterogeneity, we established pairs of standardised, debarked snags in a relatively homogeneous coniferous forest. Within each pair, one snag was fitted with an artificial hole mimicking the entrance to a woodpecker-excavated cavity, whereas the other served as a control and lacked such an entrance. This experimental design enabled us to compare bird visitation rates, species richness and behaviour between snags with artificial cavity entrances and visually similar control snags. Our aim was to determine the extent to which a single cavity entrance can act as a local attractor and influence fine-scale habitat use by forest birds. We predicted that snags with artificial cavity entrances would receive more bird visits and be visited by more species than control snags without such entrances.

## 2. Materials and methods

### 2.1. Study area and experimental design

The study was conducted in the Niepołomice Forest, a forest complex located near the city of Kraków in southern Poland, Central Europe. The study area is dominated by coniferous stands, with Scots pine *Pinus sylvestris* prevailing in the upper canopy and admixtures of pedunculate oak *Quercus robur*, silver birch *Betula pendula* and black alder *Alnus glutinosa* in the middle and lower canopy layers. The study sites were located in relatively homogeneous, predominantly single-storied stands in which the dominant Scots pines were of similar age (approximately 80 years).

The study was conducted at 30 pairs of plots (Fig. 1), each comprising two experimental snags: one fitted with an artificial cavity entrance (hereafter, snag with cavity) and one without an artificial cavity entrance (hereafter, snag without cavity). At each plot, a debarked spruce snag approximately 2.5 m tall and 20 cm in diameter was installed (Fig. 1). Each snag was embedded approximately 0.5 m into the ground to ensure stability, and its top was shaped to resemble that of a naturally broken dead tree. Within each pair, an artificial cavity entrance was created in one snag, whereas the other served as a control. The entrance was designed to mimic the entrance of a woodpecker-excavated cavity: a circular hole 5 cm in diameter was drilled 1.5 m above the ground and oriented towards the southeast. The drilled holes were approximately 10 cm deep, with their interiors inclined at an angle of approximately 25° to achieve a modest increase in depth. To prevent birds from using the artificial cavities as nesting sites during the study, the holes were made sufficiently shallow to preclude nesting. This measure was intended to minimise potential bias in observations arising from increased movement and vocalisation associated with territorial and breeding activities.

**Fig. 1.**
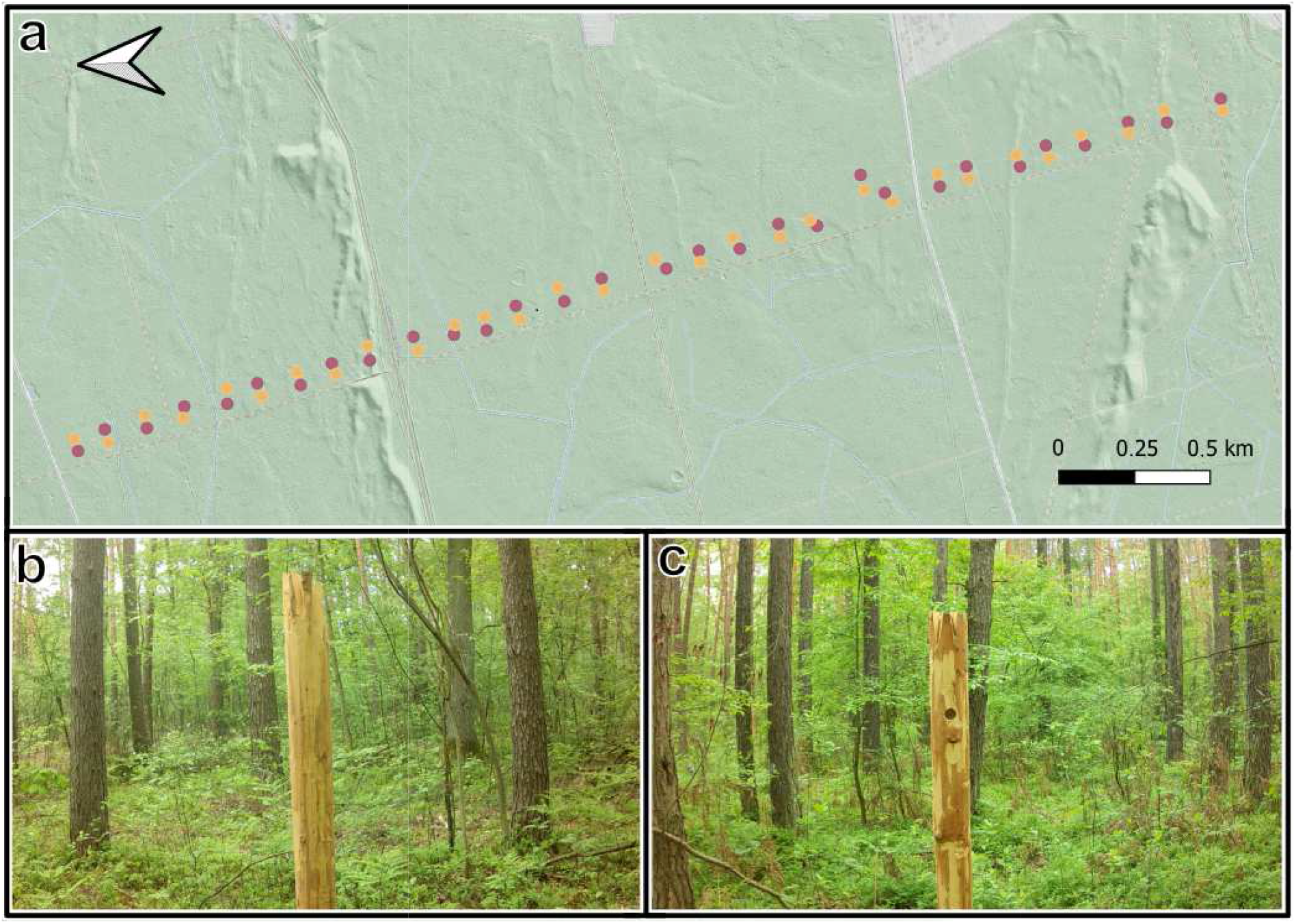
Study area and experimental design: (a) location of the 30 pairs of experimental snags in the Niepołomice Forest (southern Poland, Central Europe); (b) snag without an artificial cavity entrance (yellow); and (c) snag with an artificial cavity entrance (purple).

Within each pair, the mean distance between the two experimental snags was 34.3 ± 4.8 SD m (range: 24–43 m), whereas the mean distance between neighbouring plot pairs was 137.7 ± 29.9 SD m (range: 100–209 m). To avoid systematic spatial bias, the positions of cavity and control snags were alternated within each pair (Fig. 1). All snags were located at least 60 m from the nearest forest gravel road, which was used only occasionally by forest workers and hikers. The experimental snags were installed between 14 and 17 March 2025, and camera traps recorded bird activity over a two-month period, from 24 April to 27 June 2025.

### 2.2. Field methods

Vegetation surveys were conducted in September 2025 within circular subplots with a radius of 10 m centred on each snag. The following variables were measured within each subplot: canopy closure in the upper canopy layer, expressed as the percentage of the sky obscured by tree crowns (hereafter, Canopy); canopy closure in the middle canopy layer, expressed as the percentage of the sky obscured by crowns in this layer (hereafter, Subcanopy); shrub-layer cover, defined as the percentage of the subplot covered by woody vegetation 0.5–2 m in height (hereafter, Shrub); the distance (m) from the snag to the nearest tree with a diameter at breast height of ≥ 5 cm (hereafter, Tree distance); and the number of logs (hereafter, Logs) classified as downed coarse woody debris (diameter > 7 cm and length > 2 m).

A motion-triggered camera trap (Browning Spec Ops Elite HP5) was installed at each snag. Each camera was mounted on the nearest tree and positioned to capture the upper portion of the snag and the surrounding habitat within its field of view. The mean distance between each camera and its corresponding snag was 2.87 ± 0.62 SD m (range: 1.5–4.3 m). The cameras were triggered by motion sensors and programmed to record 10-s video clips, followed by a 10-s refractory period during which motion could not trigger another recording. All video recordings were reviewed to identify birds detected by the cameras.

### 2.3. Data handling and analyses

Video recordings from camera traps (available for 29 pairs of snags because of camera malfunction in one pair) were reviewed to identify birds interacting with the experimental snags. An interaction was defined as the direct use of a snag and included the following behavioural categories: landing on and sitting on the snag surface (visitation), moving along the snag surface (exploration), sitting on top of the snag (perching) and probing the top of the snag (probing). Behaviours involving direct use of the cavity entrance were classified as cavity inspection (approaching the entrance) or cavity cleaning. In total, 443 video recordings of birds interacting with the snags were obtained, corresponding to 246 visits. A visit was defined as an event in which a bird was recorded interacting with a snag. Consecutive clips separated by less than 10 min, with no intervening recordings of other bird species, were considered part of the same visit. A single visit could include multiple behavioural categories. To quantify the frequency of cavity-associated interactions, we calculated the proportion of visits that included at least one cavity-associated behaviour (cavity inspection or cavity cleaning). We also summed the occurrences of all behavioural categories recorded at snags with cavities and calculated the proportion attributable to cavity-associated behaviours. Snag type was treated as a two-level categorical variable (snag with cavity vs snag without cavity). Habitat characteristics at each plot were described using the following variables: Canopy, Subcanopy, Shrub, Tree distance and Logs.

To investigate species-specific responses, we calculated the following metrics for each species and snag type (snag with cavity vs snag without cavity): (i) frequency of occurrence, defined as the number of experimental snag pairs in which a species was recorded interacting with a snag of the given type; and (ii) the mean number of visits per snag (± SD), defined as the number of visits involving direct interactions with the snag. Because the presence or absence of each species at snag with or without cavity within each experimental pair constituted matched binary observations, differences in frequency of occurrence between snag types were tested using an exact McNemar test for paired binary data (Eliasziw & Donner, 1991). Differences in the number of visits between snag types were assessed using the paired Wilcoxon signed-rank test. The same species-level comparisons were performed for birds recorded by camera traps without directly interacting with the experimental snags, including birds sitting, foraging or flying in the surrounding forest (background records; Table S1). Background events were defined using the same 10-min criterion as that applied to visits involving direct interactions with snags. Differences in the frequency of non-cavity-associated behaviours between snag types were tested using paired Wilcoxon signed-rank tests. We compared the total number of non-cavity-associated behaviours recorded at snags with or without cavities, and then analysed each behavioural category separately.

To investigate the effects of cavity entrance presence and habitat characteristics on bird activity, we fitted generalized linear mixed models (GLMMs). We modelled two response variables: the number of visits per snag (visitation rate) and the number of bird species recorded per snag (species richness). Visitation rate was modelled using a negative binomial error distribution, whereas species richness was modelled using a Poisson error distribution; both models used log-link functions. The initial models included snag type (snag with cavity vs snag without cavity) and habitat characteristics as fixed effects, with snag-pair identity included as a random effect. Candidate models incorporating different combinations of predictors were compared using the Akaike information criterion corrected for small sample sizes (AICc), with lower ΔAICc values indicating greater support. For predictors retained in the final models, marginal R^2^ was partitioned into fractions uniquely attributable to snag type and habitat characteristics, as well as a fraction jointly explained by both. In addition, background bird activity and species richness were compared between snag types using negative binomial and Poisson GLMMs, respectively (Table S1). Significance level was set at p < 0.05. All statistical analyses were conducted in R (R Core Team, 2023). Models were fitted using the glmmTMB package, model selection was performed using MuMIn, and marginal R² was calculated using the performance package.

## 3. Results

### 3.1. Bird assemblage at experimental snags

In total, 246 interaction events involving 13 bird species were recorded at snags with and without cavity (Table 1). Great spotted woodpecker *Dendrocopos major* and Eurasian nuthatch *Sitta europaea* were the most frequently recorded species, while marsh/willow tit *Poecile palustris/montanus*, crested tit *Lophophanes cristatus* and great tit *Parus major* also contributed substantially to the total number of interactions, particularly at snags with cavity (Table 1). The alluvial diagram shows that most interactions involved a few common primary and secondary cavity-nesting species. Species composition was broadly similar between snag types, although the relative contribution of some species was greater at snags with cavity (Fig. 2). The frequency of occurrence did not differ between snag types for any species (Table 1). The number of interactions was higher at snags with cavity for Eurasian nuthatch (p = 0.026), whereas the differences for great spotted woodpecker (p = 0.051) and marsh/willow tit (p = 0.058) approached the adopted significance level (Table 1).

**Fig. 2.**
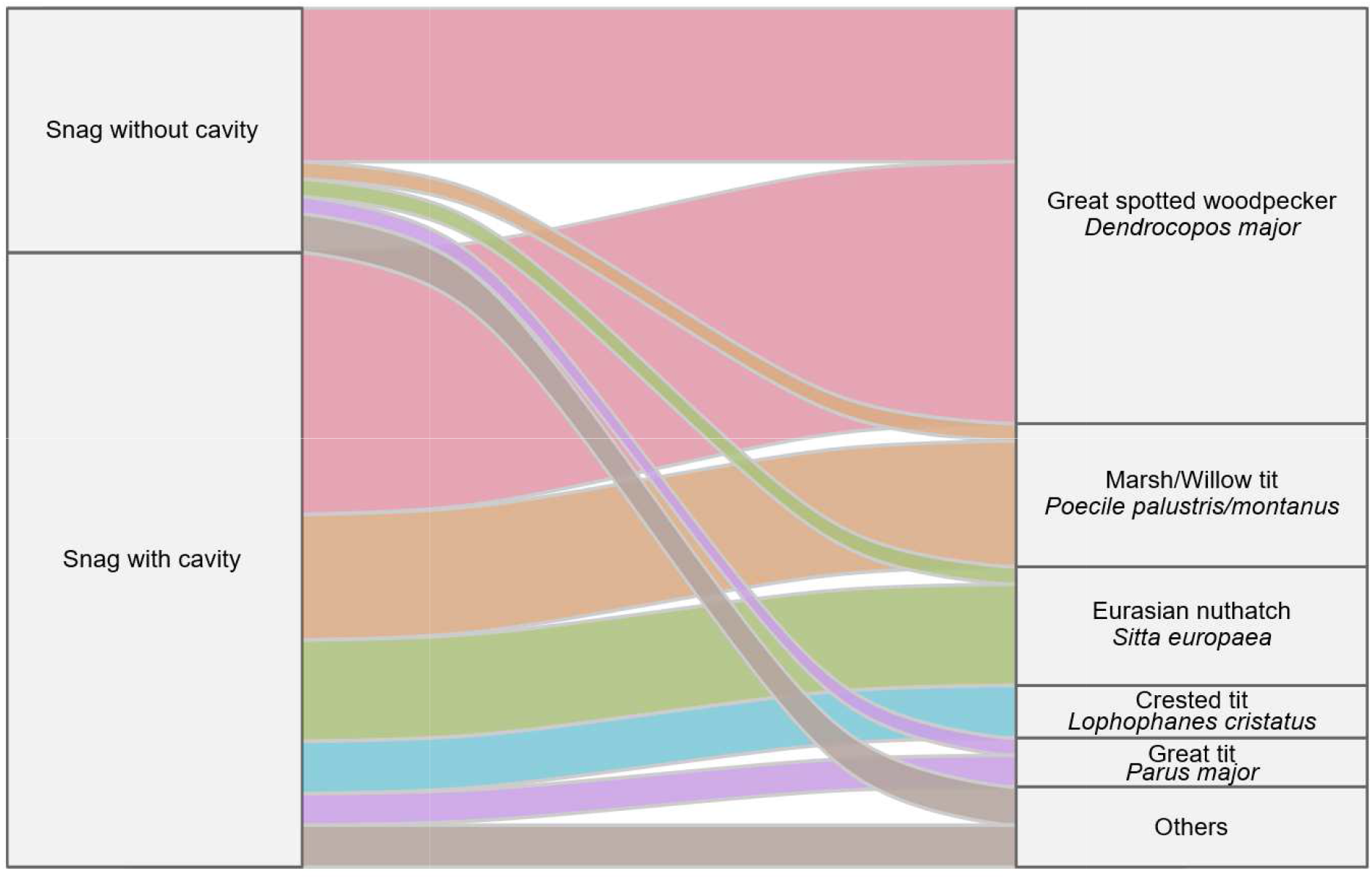
Alluvial diagram illustrating the relationship between bird species and snag type (snag with cavity vs snag without cavity) at experimental snag pairs in coniferous stands in the Niepołomice Forest (southern Poland, Central Europe). The width of each flow is proportional to the number of interaction events recorded for a given species and snag type, illustrating the relative contribution of individual species to the total number of interactions. Species with fewer than 10 interaction events are grouped under Others: common buzzard *Buteo buteo*, Ural owl *Strix uralensis*, European robin *Erithacus rubecula*, black woodpecker *Dryocopus martius*, coal tit *Periparus ater*, Eurasian jay *Garrulus glandarius*, tawny owl *Strix aluco* and spotted flycatcher *Muscicapa striata*.

**Table 1.** Frequency of occurrence (number of snags at which a given species was recorded) and mean number of interaction events per snag (± SD) for bird species recorded at experimental snag pairs (snag with cavity vs snag without cavity) in the Niepołomice Forest (southern Poland, Central Europe). Differences in frequency of occurrence between snag types were tested using the exact McNemar χ^2^ test and differences in the number of interaction events were assessed using paired Wilcoxon signed-rank tests.

| Species | Scientific name | Frequency |  |  |  | Number of interactions |  |  |  |  |  |
|---|---|---|---|---|---|---|---|---|---|---|---|
|  |  | Snag without cavity | Snag with cavity | χ2 | p | Snag without cavity |  | Snag with cavity |  | W | p |
|  |  |  |  |  |  | Mean | SD | Mean | SD |  |  |
| Great spotted woodpecker | *Dendrocopos major* | 17 | 23 | 2.1 | 0.149 | 1.5 | 1.6 | 2.6 | 2.5 | 218.0 | 0.051 |
| Eurasian nuthatch | *Sitta europaea* | 4 | 11 | 3.3 | 0.070 | 0.2 | 0.5 | 1.0 | 1.8 | 67.5 | 0.026 |
| Great tit | *Parus major* | 5 | 7 | 0.1 | 0.724 | 0.2 | 0.4 | 0.3 | 0.6 | 25.5 | 0.305 |
| Common buzzard | *Buteo buteo* | 4 | 5 | 0.0 | 1.000 | 0.1 | 0.4 | 0.2 | 0.4 | 16.0 | 0.777 |
| Marsh/Willow tit | *Poecile palustris/montanus* | 3 | 5 | 0.5 | 0.480 | 0.2 | 0.5 | 1.2 | 4.6 | 15.0 | 0.058 |
| Eurasian jay | *Garrulus glandarius* | 2 | 2 | 0.0 | 1.000 | 0.1 | 0.3 | 0.1 | 0.3 | 5.0 | 1.000 |
| Tawny owl | *Strix aluco* | 2 | 0 | 0.5 | 0.480 | 0.1 | 0.3 | 0.0 | 0.0 | 0.0 | 0.346 |
| Ural owl | *Strix uralensis* | 1 | 1 | 0.0 | 1.000 | 0.0 | 0.2 | 0.0 | 0.2 | 1.5 | 1.000 |
| European robin | *Erithacus rubecula* | 0 | 2 | 0.5 | 0.480 | 0.0 | 0.0 | 0.1 | 0.3 | 3.0 | 0.346 |
| Crested tit | *Lophophanes cristatus* | 0 | 2 | 0.5 | 0.480 | 0.0 | 0.0 | 0.5 | 2.4 | 3.0 | 0.371 |
| Coal tit | *Periparus ater* | 0 | 2 | 0.5 | 0.480 | 0.0 | 0.0 | 0.1 | 0.3 | 3.0 | 0.346 |
| Black woodpecker | *Dryocopus martius* | 1 | 0 | 0.0 | 1.000 | 0.0 | 0.2 | 0.0 | 0.0 | 0.0 | 1.000 |
| Spotted flycatcher | *Muscicapa striata* | 1 | 0 | 0.0 | 1.000 | 0.0 | 0.2 | 0.0 | 0.0 | 0.0 | 1.000 |

In total, 100 interaction events involving 20 bird species were recorded in the background (Table S1). Background bird activity did not differ between snag types (p = 0.595), with an estimated 1.5 (95% CI: 1.0–2.1) events per snag without a cavity and 1.6 (95% CI: 1.1–2.4) events per snag with a cavity. Similarly, background species richness did not differ between snag types (p = 0.739), with an estimated 1.3 species (95% CI: 0.9–1.9) per snag without a cavity and 1.4 species (95% CI: 1.0–2.0) per snag with a cavity.

### 3.2. Effects of cavity presence on visitation rate and species richness

The presence of a cavity and greater canopy closure of the dominant tree layer were positively associated with bird visitation rate (Table 2, Fig. 3a). The estimated mean visitation rate was 2.1 per snag without a cavity (95% CI: 1.4–3.2) and 4.5 per snag with a cavity (95% CI: 3.1–6.5). Visitation rate also increased with canopy closure, indicating greater bird activity at snags located in stands with a more closed canopy (Table 2). The model including both snag type and canopy closure had the lowest AICc among the candidate models (Table 3).

**Fig. 3.**
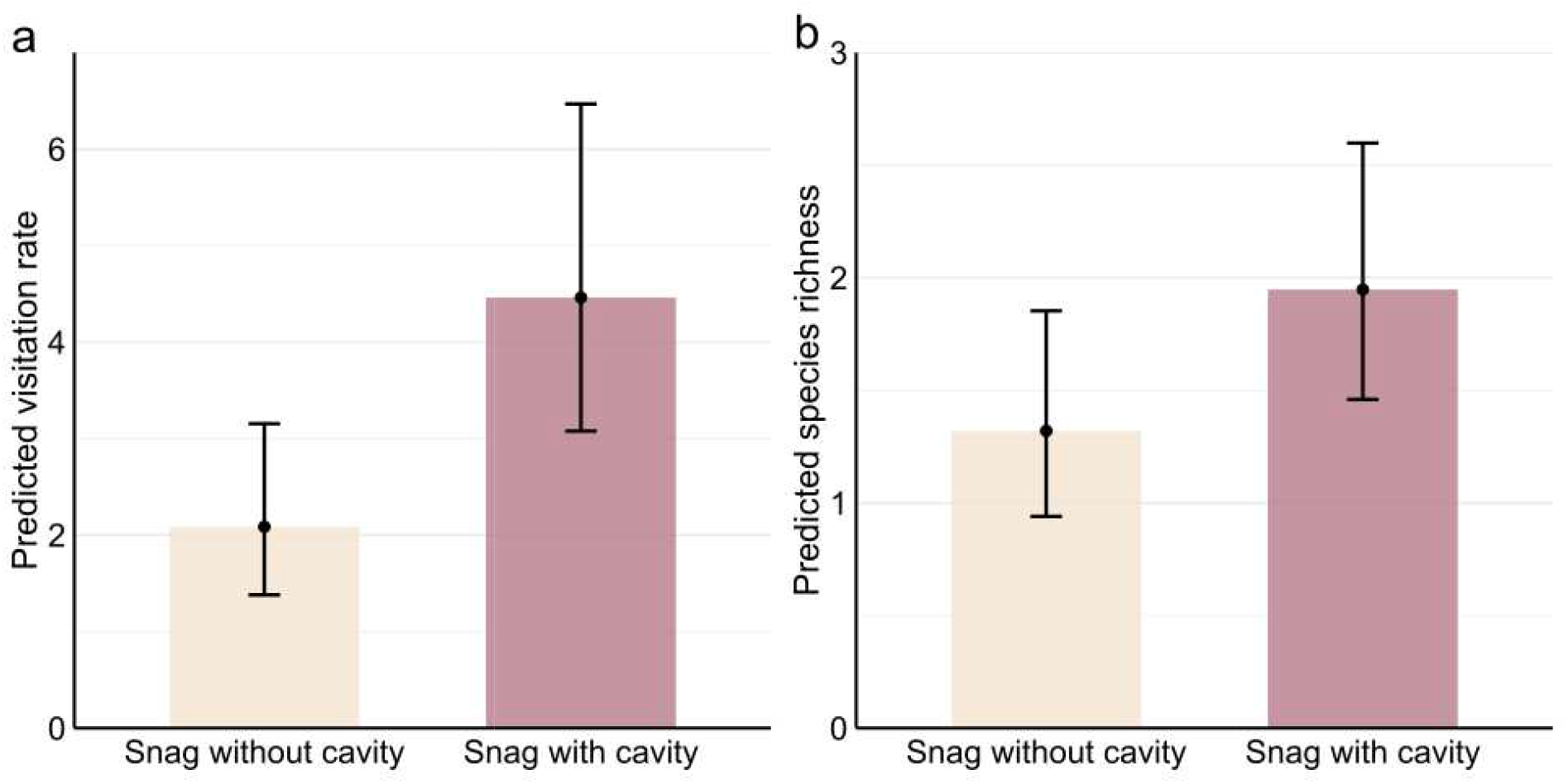
Predicted mean visitation rate (a) and species richness (b), with 95% confidence intervals, for snags without cavity and snags with cavity recorded at experimental snag pairs located in coniferous stands in the Niepołomice Forest (southern Poland, Central Europe).

**Table 2.** Generalized linear mixed models examining the relationships of bird visitation rate and species richness with snag type (snag with cavity vs snag without cavity) and habitat characteristics: canopy closure of the dominant tree layer (Canopy), canopy closure of the middle canopy layer (Subcanopy), shrub-layer cover (Shrub), distance to the nearest tree (Tree distance), and number of logs (Logs). The study was conducted at experimental snag pairs in coniferous stands in the Niepołomice Forest (southern Poland, Central Europe).

| Variable | Starting model |  |  |  | Final model |  |  |  |
|---|---|---|---|---|---|---|---|---|
|  | Estimate | SE | Z | p | Estimate | SE | Z | p |
| Visitation rate |  |  |  |  |  |  |  |  |
| Intercept | -1.13 | 1.09 | -1.0 | 0.302 | -0.89 | 0.72 | -1.3 | 0.212 |
| Snag type (cavity) | 0.76 | 0.23 | 3.3 | 0.001 | 0.76 | 0.23 | 3.4 | 0.001 |
| Canopy | 0.04 | 0.02 | 2.5 | 0.013 | 0.04 | 0.02 | 2.4 | 0.016 |
| Subcanopy | 0.01 | 0.01 | 0.8 | 0.448 |  |  |  |  |
| Shrub | 0.00 | 0.02 | 0.2 | 0.857 |  |  |  |  |
| Tree distance | -0.03 | 0.24 | -0.1 | 0.891 |  |  |  |  |
| Logs | -0.02 | 0.06 | -0.3 | 0.781 |  |  |  |  |
| Species richness |  |  |  |  |  |  |  |  |
| Intercept | -0.58 | 0.83 | -0.7 | 0.480 | -0.55 | 0.54 | -1.0 | 0.311 |
| Snag type (cavity) | 0.39 | 0.21 | 1.9 | 0.063 | 0.39 | 0.20 | 1.9 | 0.057 |
| Canopy | 0.02 | 0.01 | 1.7 | 0.097 | 0.02 | 0.01 | 1.7 | 0.098 |
| Subcanopy | 0.01 | 0.01 | 1.1 | 0.270 |  |  |  |  |
| Shrub | 0.01 | 0.01 | 0.7 | 0.459 |  |  |  |  |
| Tree distance | -0.10 | 0.18 | -0.6 | 0.556 |  |  |  |  |
| Logs | -0.06 | 0.05 | -1.2 | 0.239 |  |  |  |  |

**Table 3.**
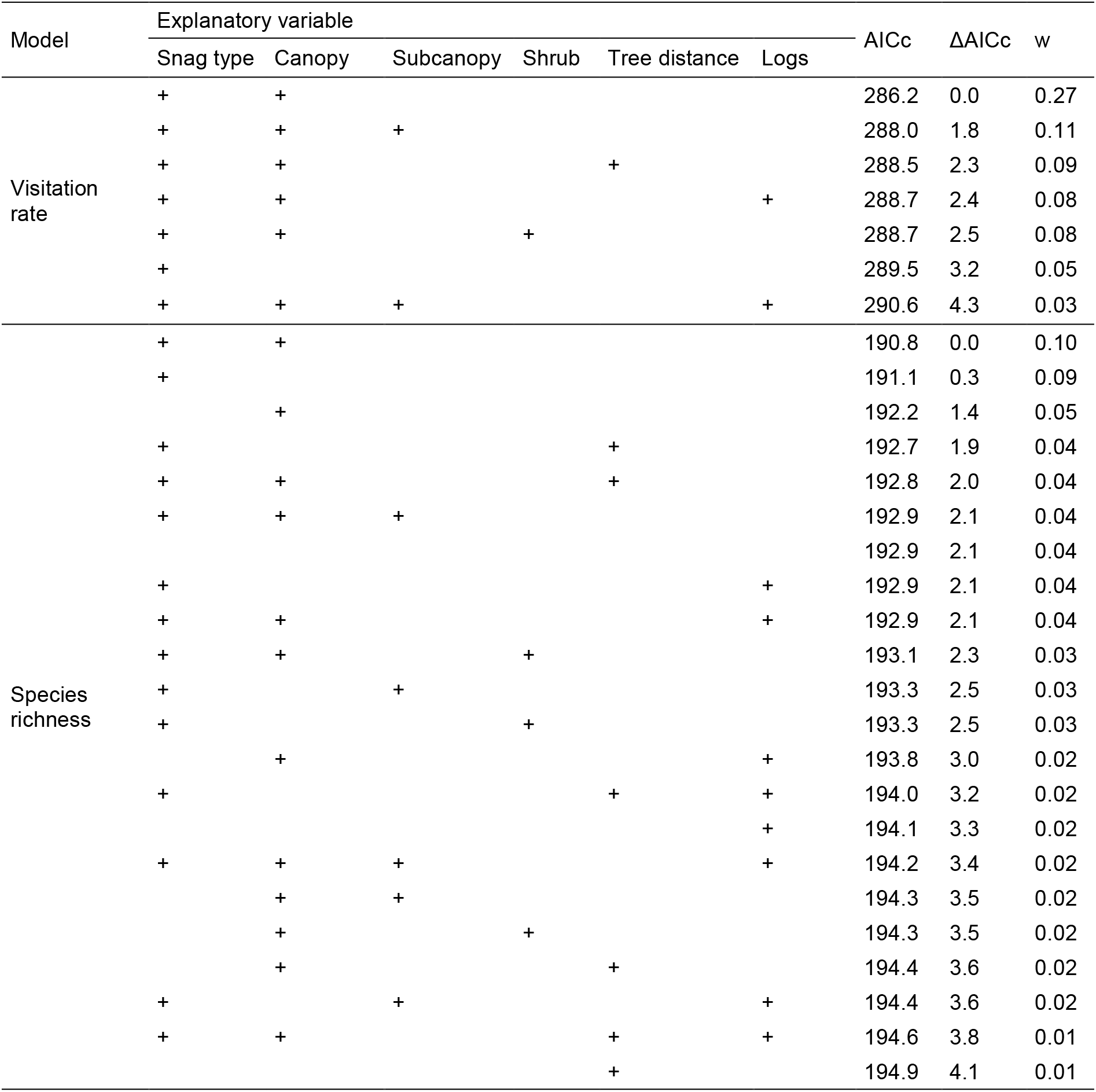
Candidate models explaining variation in bird visitation rate and species richness at experimental snag pairs (snag with cavity vs snag without cavity) in coniferous stands in the Niepołomice Forest (southern Poland, Central Europe). The candidate predictors included snag type (presence or absence of an artificial cavity entrance) and habitat characteristics: canopy closure of the dominant tree layer (Canopy), canopy closure of the middle canopy layer (Subcanopy), shrub-layer cover (Shrub), distance to the nearest tree (Tree distance), and number of logs (Logs). Models are ranked by increasing values of the Akaike information criterion corrected for small sample sizes (AICc). For each model, the difference in AICc relative to the top-ranked model (ΔAICc) and the Akaike weight (w) are reported.

Species richness was higher (approached the adopted significance level) at snags with cavities than at snags without cavities (Table 2, Fig. 3b). The estimated mean species richness was 1.3 per snag without a cavity (95% CI: 0.9–1.9) and 2.0 per snag with a cavity (95% CI: 1.5–2.6). The model including both cavity presence and canopy closure of the dominant tree layer was the highest-ranked model, although the model including cavity presence alone received comparable support (Table 3).

Partitioning of marginal R^2^ indicated that both cavity presence and canopy closure of the dominant tree layer contributed to explaining variation in bird responses at experimental snags (Fig. 4). For visitation rate, the model explained 27.5% of the variance, with snag type and canopy closure uniquely accounting for 11.3% and 11.1%, respectively, and a further 5.0% jointly explained by both predictors (Fig. 4a). For species richness, the model explained 12.5% of the variance, with snag type and canopy closure uniquely accounting for 6.4% and 5.5%, respectively, and a further 0.5% jointly explained by both predictors (Fig. 4b).

**Fig. 4.**
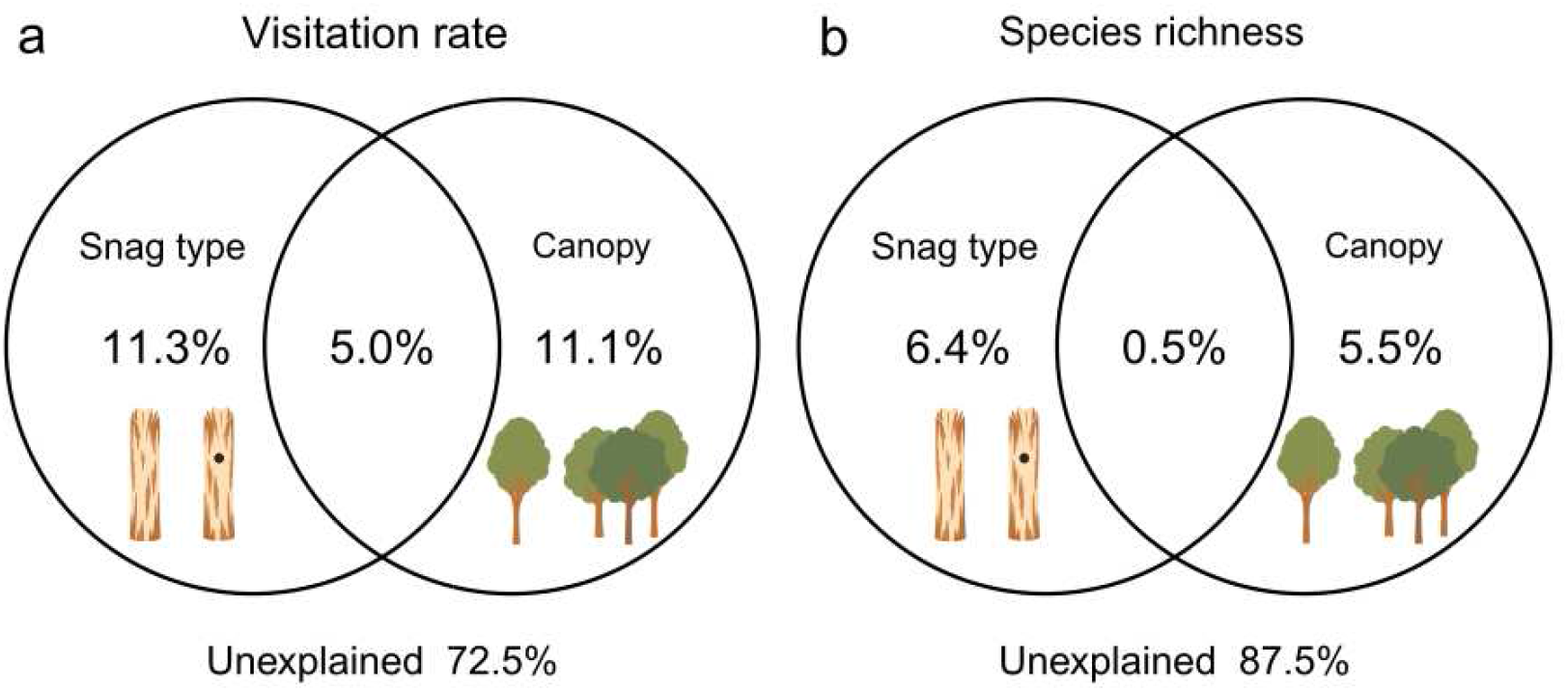
Partitioning of marginal R^2^ showing the contributions of snag type (snag with cavity vs snag without cavity) and habitat characteristics to variation in (a) bird visitation rate and (b) bird species richness at experimental snag pairs in coniferous stands in the Niepołomice Forest (southern Poland, Central Europe). For both response variables, the unique fractions of variance explained by snag type and canopy closure of the dominant tree layer (Canopy), as well as the fraction jointly explained by both predictors, are shown.

### 3.3. Species and behaviours at experimental snags

Great spotted woodpecker, marsh/willow tit and Eurasian nuthatch accounted for most interactions involving direct use of the experimental snags, while the crested tit and great tit contributed additional records (Fig. 5). The presence of a cavity entrance provided additional opportunities for snag use, resulting in two additional behavioural categories: cavity inspection and cavity cleaning. Cavity-associated behaviours occurred in 73.3% of all interaction events recorded at snags with cavities and accounted for 42.3% of all behavioural occurrences recorded at these snags (Fig. 5). Species classified as Others, primarily raptors (common buzzard *Buteo buteo*) and owls (tawny owl *Strix aluco* and Ural owl *Strix uralensis*), used the snags as perches (Fig. 5). Overall, non-cavity-associated behaviours, i.e. visitation, exploration, perching and probing, were recorded more frequently at snags with cavities than at snags without cavities, with the difference approaching the adopted significance level (W = 264, p = 0.073; Fig. 5). When individual behavioural categories were analysed separately, exploration occurred more frequently at snags with cavities (W = 187, p = 0.049), whereas the frequencies of the other non-cavity-associated behaviours, i.e. visitation, perching and probing, did not differ between snag types (all p ≥ 0.129).

**Fig. 5.**
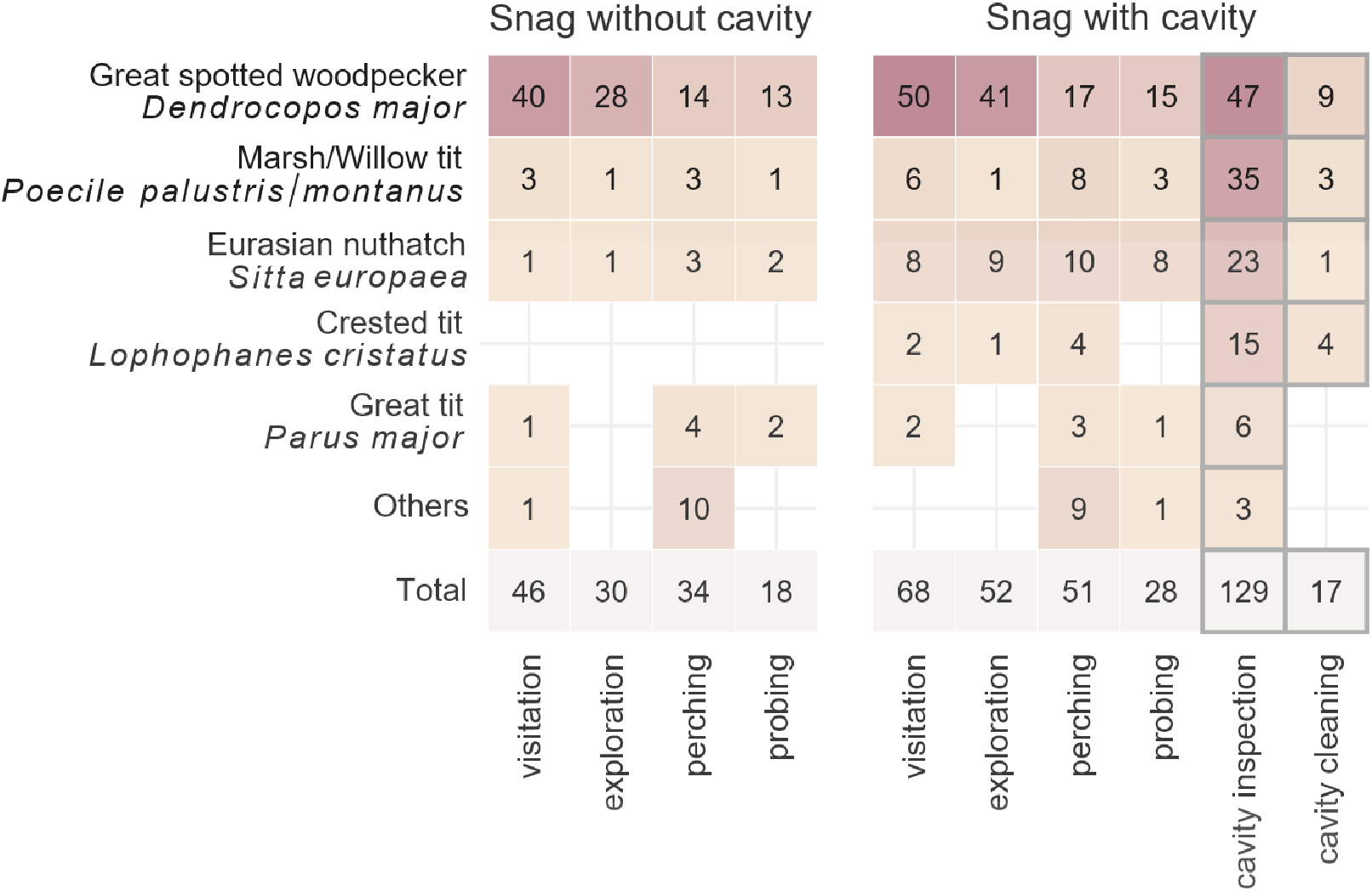
Number and types of interactions recorded for bird species at experimental snags with cavities and snags without cavities in coniferous stands in the Niepołomice Forest (southern Poland, Central Europe). Cell colour intensity is proportional to the number of recorded interactions. Grey outlines indicate cavity-associated behaviours. Species with fewer than 10 recorded interactions are grouped under Others: common buzzard *Buteo buteo*, Ural owl *Strix uralensis*, European robin *Erithacus rubecula*, black woodpecker *Dryocopus martius*, coal tit *Periparus ater*, Eurasian jay *Garrulus glandarius*, tawny owl *Strix aluco* and spotted flycatcher *Muscicapa striata*.

## 4. Discussion

### 4.1. Cavity presence as a local attractor for forest birds

Our results demonstrate that the presence of a single cavity entrance in a snag influences bird activity and species richness in a structurally homogeneous forest. Snags with an artificial opening mimicking a woodpecker-excavated cavity entrance received 114% more visits and had 48% higher estimated species richness than control snags without cavity entrances. Because the snags were standardised in size and shape and their positions were independent from habitat context, our findings suggest that the presence of a cavity can influence fine-scale patterns of habitat use, supporting the hypothesis that cavity entrances act as local attractors for forest birds.

Tree cavities provide potential sites for breeding, resting and foraging (Aitken & Martin, 2007; Tarbill et al., 2015). Consequently, complex networks of interactions may develop around cavity entrances involving predators, competitors and potential mates (Drever et al., 2008; Martin, 2015; Kotowska et al., 2020; Van Der Hoek et al., 2020). Our results suggest that cavity entrances may serve as hotspots for intra-and interspecific interactions, potentially by providing cues that convey information about the presence or previous activity of other organisms. These findings contribute to our understanding of the nest-web concept, which describes a network of ecological relationships linking primary cavity excavators, the cavities they create and cavity-dependent species (Martin & Eadie, 1999; Martin et al., 2004; Martin, 2015; Cockle et al., 2019). Whereas previous studies have largely focused on relationships between cavity availability and community-level patterns in the abundance and diversity of secondary cavity users, our experimental results indicate that even a single cavity entrance can increase visitation rates and the number of bird species recorded at its immediate surroundings.

Species closely associated with tree cavities, particularly great spotted woodpecker, Eurasian nuthatch and marsh/willow tit, accounted for most recorded interactions. These species use tree cavities as nesting and roosting sites (Wesołowski, 1996; Krištín et al., 2001; Mazgajski, 2002; Maziarz et al., 2015; Parry & Broughton, 2018). Cavity-nesting birds are known to inspect cavities and use social and environmental cues to assess their suitability (Mazgajski, 2007; Podofillini et al., 2018). In our study, birds frequently approached the artificial cavity entrances and, in some cases, probed or manipulated woody material that may persist inside the openings. Such behaviours are consistent with cavity inspection and prospecting for potential nesting or roosting sites and may allow birds to gather information about cavity suitability and previous or current occupancy. Moreover, exploration, defined as movement along the snag surface, occurred more frequently at snags with cavity entrances than at control snags without cavities, suggesting that the presence of a cavity entrance influenced bird behaviour. This increased exploratory activity may reflect more intensive investigation of snags bearing cavity entrances (Mazgajski, 2007; Podofillini et al., 2018). Alternatively, birds may have explored snags with cavities more frequently because cavity-bearing trees often host other tree-related microhabitats that provide potential food resources (Gutzat & Dormann, 2018; Wesołowski & Martin, 2018). In contrast, large-bodied species, such as common buzzard and owls, used both snag types as perching sites, with no recorded interactions with the cavity entrances. These behavioural differences suggest that responses to the experimental snags may vary according to species-specific associations with tree cavities, with cavity-associated species showing both direct interactions with the entrances and increased exploratory behaviour.

### 4.2. Cavity presence and habitat characteristics

Our study shows that habitat characteristics, together with cavity presence, explain variation in visitation rates and species richness at experimental snags. Canopy closure in the upper layer was associated with more visits and a higher number of bird species recorded, suggesting that birds responding to cavity entrances may preferentially use habitats with greater canopy closure, possibly because these habitats provide greater concealment. This pattern contrasts with previous studies reporting higher bird species richness and abundance in small-scale canopy gaps than in adjacent closed-canopy forests (Fuller, 2000; Lewandowski et al., 2021). More open, sun-exposed patches, together with the availability of coarse woody debris that provides substrate for numerous invertebrates, can support greater food availability for insectivorous birds and thereby increase local bird richness and abundance (Moorman et al., 2007; Perry et al., 2018; Przepióra et al., 2020). However, these studies of bird assemblages associated with canopy gaps relied on breeding bird surveys, which primarily detect vocalising and displaying individuals, whereas our camera-trap-based experimental design recorded birds directly interacting with snags for various purposes. The positive association between canopy closure and visitation rates may therefore partly reflect the behaviour of cavity-associated birds prospecting for potential nesting or roosting sites. Vegetation cover around cavities may also provide greater concealment from predators, potentially reducing nest predation risk for some cavity-nesting birds (Fisher & Wiebe, 2006; Berl et al., 2014).

Closed-canopy forests provide buffered microclimatic conditions (De Frenne et al., 2019), which may influence arthropod distribution and activity (Sallé et al., 2021). Folivorous caterpillars are an important food resource for birds, particularly during the breeding season, including for cavity-nesting species (Wesołowski & Neubauer, 2017; Wesołowski et al., 2019). Moreover, foraging birds may select habitat patches by balancing food availability against proximity to protective cover and the associated risk of predation (Walther & Gosler, 2001; Villén-Pérez et al., 2013). Greater canopy closure around experimental snags may therefore enable birds to move through the habitat with reduced exposure to predators. This may be particularly important during the breeding season, when frequent food provisioning increases parental activity around nests and may consequently increase the risk of nest failure due to predation (Martin et al., 2000; Yoon et al., 2016). Because a substantial proportion of the variation in both visitation rates and species richness remained unexplained, other habitat-, social-, and community-related factors, such as food availability and the presence of competitors, predators or potential mates, may also influence bird activity around cavity entrances. Nevertheless, cavity presence alone explained a proportion of the variation in visitation rates and species richness comparable to that explained by canopy closure, a habitat characteristic frequently identified as an important determinant of bird community composition and abundance.

### 4.3. Limitations and implications

Our study examined a relatively simple cavity design, with a standardised entrance height and orientation, and covered only a limited period within a single breeding season. Variation in cavity dimensions, depth and entrance height may elicit different levels of interest or behavioural responses from birds, as these characteristics are known to influence cavity use and nest-site selection (Wesołowski, 2003; Wesołowski & Rowiński, 2004; Di Sallo & Cockle, 2022). Moreover, our experiment was conducted in managed forest stands, where tree cavities are less abundant than in unmanaged forests (Remm & Lõhmus, 2011; Walankiewicz et al., 2014). Responses to newly available cavities may therefore depend on overall cavity availability, with individual cavities potentially having greater ecological importance where this resource is scarce (Cockle et al., 2011; Wiebe, 2011). In our experimental design, the snags were installed several weeks before camera-trap monitoring began. Birds may therefore have become familiar with the snags themselves, reducing the likelihood that the observed activity reflected an initial response to novelty, such as neophilic attraction to or neophobic avoidance of a newly introduced vertical structure (Greenberg & Mettke-Hofmann, 2001; Tryjanowski et al., 2016). The observed differences between snags with and without cavity entrances therefore suggest that the presence of a cavity entrance, rather than the mere presence of a snag, was associated with increased bird activity. Despite these limitations, our paired experimental design, which used standardised snags differing only in the presence or absence of a cavity entrance within a relatively homogeneous habitat, provides evidence that a single cavity entrance can act as a local attractor for forest birds.

## 5. Conclusions

By demonstrating that a single cavity can act as a fine-scale attractor for birds, our study extends current understanding of nest-web functioning, in which cavity availability and primary cavity excavators shape interactions within communities of cavity-using species (Martin & Eadie, 1999; Cockle et al., 2011; Van Der Hoek et al., 2020). Our results indicate that cavities may influence bird behaviour even when they are not associated with active nesting, and that a simple entrance hole, in the absence of breeding-associated cues such as calls, faeces or prey remains, may be sufficient to elicit cavity inspection and repeated visits by cavity-using species. Thus, individual cavities may function not only as access points to potential nesting or roosting sites but also as focal points of bird activity at a fine spatial scale. These findings highlight the potential for small-scale habitat features to create behavioural hotspots within forest landscapes (Maciorowski et al., 2021), potentially concentrating inter-and intraspecific interactions, the exchange of social information, and predator–prey encounters around structures such as cavities. Conservation and management measures that retain old trees and cavity-bearing trees, particularly in managed forests where cavities may be scarce, could therefore benefit cavity-using communities beyond the direct provision of nesting sites (Winter & Möller, 2008; Gutzat & Dormann, 2018). More broadly, our findings suggest that the ecological significance of cavities may extend beyond their direct use as nesting sites, as cavities that are not associated with active breeding may nevertheless serve as local centres of bird activity.

## Acknowledgments

We thank Michał Wieciech from the Niepołomice State Forest District for support in conducting the study.

## Funding declaration

This research was funded in part by the National Science Centre, Poland (No. 2021/41/B/NZ8/03456). AW was supported by the funds for students of the Doctoral School of the University of Agriculture in Kraków.

## Authorship contribution statement

**Alicja Wolska:** Data curation, Formal analysis, Investigation, Resources, Visualization, Writing - original draft, Writing - review & editing; Marcin Trybała: Methodology, Resources; **Michał Ciach:** Conceptualization, Formal analysis, Funding acquisition, Investigation, Methodology, Supervision, Writing - original draft, Writing - review & editing.

## Ethical statement

The study was performed in accordance with Polish law.

## Conflict of interest

The authors declare that they have no known competing financial interests or personal relationships that could have appeared to influence the work reported in this paper.

## Data availability

The data used in this study is available on request from the corresponding author.

---

**Table S1.** Frequency of occurrence (number of plots on which a given species was recorded) and mean ± SD number of background events per plot for bird species recorded in the surroundings of experimental snag pairs (snag with cavity vs snag without cavity) in the Niepołomice Forest (southern Poland, Central Europe). Species recorded interacting with experimental snags are listed first, in the same order as in Table 1, including those not recorded in the background (indicated by zero). Species recorded exclusively in the background are presented separately.

| Species | Scientific name | Frequency |  |  |  | Number of events |  |  |  |  |  |
|---|---|---|---|---|---|---|---|---|---|---|---|
|  |  | Plot with snag without cavity | Plot with snag with cavity | χ2 | p | Plot with snag without cavity |  | Plot with snag with cavity |  | W | p |
|  |  |  |  |  |  | Mean | SD | Mean | SD |  |  |
| Species recorded on snags |  |  |  |  |  |  |  |  |  |  |  |
| Great spotted woodpecker | *Dendrocopos major* | 8 | 7 | 0.0 | 1.000 | 0.34 | 0.61 | 0.31 | 0.60 | 25.0 | 0.824 |
| Eurasian nuthatch | *Sitta europaea* | 3 | 4 | 0.0 | 1.000 | 0.10 | 0.31 | 0.17 | 0.47 | 17.5 | 0.588 |
| Great tit | *Parus major* | 8 | 8 | 0.0 | 1.000 | 0.38 | 0.73 | 0.34 | 0.61 | 52.0 | 1.000 |
| Common buzzard | *Buteo buteo* | 3 | 1 | 0.3 | 0.617 | 0.10 | 0.31 | 0.03 | 0.19 | 2.5 | 0.424 |
| Marsh/Willow tit | *Poecile palustris/montanus* | 1 | 1 | 0.0 | 1.000 | 0.03 | 0.19 | 0.03 | 0.19 | 1.5 | 1.000 |
| Eurasian jay | *Garrulus glandarius* | 2 | 5 | 0.6 | 0.450 | 0.07 | 0.26 | 0.17 | 0.38 | 20.0 | 0.299 |
| Tawny owl | *Strix aluco* | 0 | 2 | 0.5 | 0.480 | 0.00 | 0.00 | 0.10 | 0.41 | 3.0 | 0.371 |
| Ural owl | *Strix uralensis* | 1 | 0 | 0.0 | 1.000 | 0.10 | 0.56 | 0.00 | 0.00 | 0.0 | 1.000 |
| European robin | *Erithacus rubecula* | 0 | 4 | 2.3 | 0.134 | 0.00 | 0.00 | 0.14 | 0.35 | 10.0 | 0.072 |
| Crested tit | *Lophophanes cristatus* | 1 | 0 | 0.0 | 1.000 | 0.03 | 0.19 | 0.00 | 0.00 | 0.0 | 1.000 |
| Coal tit | *Periparus ater* | 0 | 0 |  |  |  |  |  |  |  |  |
| Black woodpecker | *Dryocopus martius* | 2 | 0 | 0.5 | 0.480 | 0.07 | 0.26 | 0.00 | 0.00 | 0.0 | 0.346 |
| Spotted flycatcher | *Muscicapa striata* | 2 | 0 | 0.5 | 0.480 | 0.07 | 0.26 | 0.00 | 0.00 | 0.0 | 0.346 |
| Species recorded exclusively in the background |  |  |  |  |  |  |  |  |  |  |  |
| Eurasian blackbird | *Turdus merula* | 2 | 2 | 0.0 | 1.000 | 0.07 | 0.26 | 0.10 | 0.41 | 6.0 | 0.850 |
| Unidentified passerine |  | 2 | 2 | 0.0 | 1.000 | 0.07 | 0.26 | 0.10 | 0.41 | 6.0 | 0.850 |
| Common chaffinch | *Fringilla coelebs* | 1 | 2 | 0.0 | 1.000 | 0.03 | 0.19 | 0.07 | 0.26 | 4.0 | 0.773 |
| Collared flycatcher | *Ficedula albicollis* | 1 | 1 | 0.0 | 1.000 | 0.03 | 0.19 | 0.03 | 0.19 | 1.5 | 1.000 |
| Eurasian blue tit | *Cyanistes caeruleus* | 1 | 1 | 0.0 | 1.000 | 0.03 | 0.19 | 0.03 | 0.19 | 0.0 | 1.000 |
| Hawfinch | *Coccothraustes coccothraustes* | 1 | 0 | 0.0 | 1.000 | 0.03 | 0.19 | 0.00 | 0.00 | 0.0 | 1.000 |
| Eurasian blackcap | *Sylvia atricapilla* | 0 | 1 | 0.0 | 1.000 | 0.00 | 0.00 | 0.03 | 0.19 | 1.0 | 1.000 |
| Common crane | *Grus grus* | 0 | 1 | 0.0 | 1.000 | 0.00 | 0.00 | 0.17 | 0.93 | 1.0 | 1.000 |

## Notes

### Competing Interest Statement

The authors have declared no competing interest.

